# Influence of insulin sensitivity on food cue evoked functional brain connectivity in children

**DOI:** 10.1101/2024.02.12.579924

**Authors:** Lorenzo Semeia, Ralf Veit, Sixiu Zhao, Shan Luo, Brendan Angelo, Andreas L. Birkenfeld, Hubert Preissl, Anny H. Xiang, Stephanie Kullmann, Kathleen A. Page

## Abstract

**Objective:** Insulin resistance during childhood is a risk factor for developing type 2 diabetes and other health problems later in life. Studies in adults have shown that insulin resistance affects regional and network activity in the brain which are vital for behavior, e.g. ingestion and metabolic control. To date, no study has investigated whether brain responses to food cues in children are associated with peripheral insulin sensitivity.

**Methods:** We included 53 children (36 girls) between the age of 7-11 years, who underwent an oral Glucose Tolerance Test (oGTT) to estimate peripheral insulin sensitivity (ISI). Brain responses were measured using functional magnetic resonance imaging (fMRI) before and after glucose ingestion. We compared food-cue task-based activity and functional connectivity (FC) between children with low and high ISI, adjusted for age and BMIz.

**Results:** Independent of prandial state (i.e., glucose ingestion), children with lower ISI showed higher FC between the anterior insula and caudate and lower FC between the posterior insula and mid temporal cortex than children with higher ISI. Sex differences were found based on prandial state and peripheral insulin sensitivity in the insular FC. No differences were found on whole-brain food-cue reactivity.

**Conclusions:** Children with low peripheral insulin sensitivity showed differences in food cue evoked response particularly in insula functional connectivity. These differences might influence eating behavior and future risk of developing diabetes.

## 1. Introduction

The prevalence of overweight and obesity among the global population is increasing worldwide, with high rates in the Americas (61%), Europe (55%), and the eastern Mediterranean (46%) (Yatsuya et al., 2014). Unfortunately, this upward trend impacts children as well, with 18% of European children between 2-7 years old having overweight/obesity (Garrido-Miguel et al., 2019). In several European countries, the number of children with overweight/obesity peaks at around 50% in those aged 6-9 years old (Buoncristiano et al., 2021), a trend similar to that observed in the American population (Fryar, Carroll, & Ogden, 2018). Overweight and obesity in childhood are risk factors for obesity and type 2 diabetes mellitus (T2DM) in adulthood (Barton, 2012), and they have adverse effects on psychological health, cognition, brain structure and function (Brooks, Smith, & Stamoulis, 2023; Wang et al., 2019). Peripheral insulin resistance, which is a condition where insulin is not able to adequately promote glucose uptake by peripheral tissues, is considered a risk factor for the future development of obesity, type 2 diabetes and cardiovascular diseases (Kahn, Hull, & Utzschneider, 2006).

Exposure to high-calorie food in the environment may promote overconsumption of unhealthy meals through stimulation of brain areas associated with reward and motivation (Pujol et al., 2021; Stice & Burger, 2019; Stice, Figlewicz, Gosnell, Levine, & Pratt, 2013). These areas include the amygdala, anterior cingulate cortex, hippocampus, hypothalamus, dorsal and ventral striatum, insula, and prefrontal cortex (Yang, Wu, & Morys, 2021). Supporting this theory, increased neural activation in response to pictures of high-caloric foods has been observed in individuals with obesity, compared to their lean counterparts, among adults, adolescents, and children (Li et al., 2023a). Of note, neural reactivity to food cues is predictive of future weight gain and linked to food craving, which may contribute to obesity onset in both children and adults (Boswell & Kober, 2016; Stice, Yokum, Bohon, Marti, & Smolen, 2010). Functional connectivity (FC) between areas of the reward network during the presentation of food cues is also enhanced in adults with obesity, increasing the motivational value of food (Stoeckel et al., 2009) and food intake (Carnell, Benson, Pantazatos, Hirsch, & Geliebter, 2014), and decreasing self-regulation (Donofry et al., 2020a; Donofry, Stillman, & Erickson, 2020b).

The current state of satiety and postprandial hormonal responses modulate regional activity and functional connections of brain regions vital for ingestive behaviour (Capucho & Conde, 2023). For example, the increase in insulin after glucose ingestion is associated with a reduction in food cue reactivity in the insula, striatum and orbitofrontal cortex in adults (Heni et al., 2014; Kroemer et al., 2013). Overall, the brain responses to glucose ingestion are greater in children compared to adults, and they are related to overweight and obesity (Ge et al., 2021). Moreover, peripheral insulin resistance in adults is linked to stronger connectivity along the reward network after a meal, suggesting a failure of insulin to suppress rewarding signals, which may increase the risk for overeating and obesity (Ryan, Karim, Aizenstein, Helbling, & Toledo, 2018). Even though glucose uptake in the brain is independent of insulin, proper insulin signalling is essential for maintaining energy homeostasis and cognitive functions (Kullmann et al., 2020). Insulin receptors are expressed throughout the brain, including the hypothalamus, striatum, amygdala, hippocampus, insula cortex, and PFC (for a review, see Kullmann et al. (2020)). In adults, central insulin action, measured by the neural response to intranasal insulin, has been found to modulate regional activity and FC of the food reward network (e.g. (Kullmann et al., 2017; Kullmann et al., 2018; Tiedemann et al., 2017)) attributed to regulating hunger, food craving and food choice differentially in adults with normal weight or obesity. First evidence points to the development of central insulin resistance in utero, as fetal brain responses to oral glucose were found to be slower in the offspring of mothers with gestational diabetes mellitus (GDM, (Linder et al., 2015). Similarly, children exposed to GDM before the 26^th^ week of gestation fail to inhibit hypothalamic activity following glucose consumption (Page et al., 2019). These differences in glucose-induced brain responses are interpreted as a possible risk factor for the development of obesity.

However, little is known about the impact of peripheral insulin sensitivity on the brain during childhood. To this end, the present study aims to explore the relationship between peripheral insulin sensitivity index (ISI) and both reactivity and whole-brain functional connectivity (FC) during a food cue task, before and after glucose ingestion, in children aged 7-11 years old. The analysis was conducted in the the BrainChild Cohort on the effects of early-life exposures on brain and metabolic health (Page et al., 2019). Prior findings in this cohort have shown greater food-cue reactivity in brain areas involved in reward and motivation (i.e., orbitofrontal cortex, amygdala, striatum, insula) among children in the fasted state (Luo et al., 2019). In addition, prior reports in the BrainChild cohort have shown that prenatal exposure to maternal obesity or gestational diabetes mellitus is associated with alterations in the structural development of the hippocampus (Alves et al., 2020; Lynch et al., 2021), as well as greater food cue reactivity in brain reward regions and greater caloric intake (Luo et al., 2021). In the current study, we hypothesize that children with low compared to children with high insulin sensitivity will exhibit higher whole brain reactivity and higher FC to food cues as compared to non-food images. Finally, we expect a decrease in neural reactivity and FC to food cues following glucose ingestion among children with higher insulin sensitivity.

## 2. Methods

### 2.1. Participants

The current study includes MRI data of 112 healthy children aged 7 to 11 years old who were recruited from the BrainChild study (Page et al., 2019) which examines risks for diabetes and obesity in children exposed to maternal diabetes during pregnancy. The children were recruited from Kaiser Permanente Southern California (KPSC). To be included, children were healthy without any neurological or psychological disorders, right-handed, and having normal or corrected vision. After preprocessing (see section 2.4. *Images preprocessing*), MRI data of 53 children were used for further data analysis.

### 2.2. Procedure

Prior to data collection, parental consent and child assent were obtained. The Institutional Review Board of the University of Southern California and KPSC approved all procedures. The baseline study visits are comprised of two separate visits conducted on different days. During the first visit, anthropometric data were collected and an oral Glucose Tolerance Test (oGTT) was performed. The second visit involved the food cue task conducted during an fMRI scan (for details see (Luo et al., 2019)).

#### First visit: anthropometrics and OGTT

The first measurements were conducted at the Clinical Research Unit of the USC Diabetes and Obesity Research Institute in the morning after a 12-hour overnight fast. Child’s anthropometric measures were collected as previously described (Page et al., 2019). BMIz (standardized BMI, specific for age and sex standard deviation) scores were computed for children based on the Center for Disease Control (CDC) normative data (2000 CDC Growth Charts for the United States: Methods and Development, 2002). Finally, a 3-hour oGTT was performed, and children’s peripheral insulin sensitivity (ISI) was estimated using the Matsuda index (Matsuda & DeFronzo, 1999).

#### Second visit: food cue task and glucose ingestion

During the second visit, which took place at the USC Dana and David Dornsife Neuroimaging Center, children underwent MRI measurements after being trained on a mock scanner. The MRI protocol consisted of a food cue task before (fasted state) and ∼15 minutes after the glucose drink. MRI scans were conducted in the morning between 8 and 10 am, after a 12-hour overnight fast. The food cue task is described in detail in (Luo et al., 2019)). In a randomized order, the participants were presented with 12 pictures of high-caloric food (F, high-caloric food such as French fries or pancakes) and 12 pictures of non-food pictures (NF, such as books and rulers) and were asked to attentively watch these pictures. Each block consisted of three pictures from one of the two stimulus types (F vs. NF), which were presented in a random order. Each image was displayed for 4 seconds and followed by a 1-second interstimulus interval. The total duration of the task was 196 seconds. After the first MRI measurement, participants consumed a glucose drink (1.75 g/kg of body weight, max 75g) to match the design of the OGTT test performed on a separate day. About 15 minutes after the glucose ingestion, participants underwent a second food cue task using the same procedure as before. The picture set in the two food cue tasks was the same, with the order of the images randomized both within and between blocks.

### 2.3. MRI data acquisition

The imaging was conducted using a Siemens MAGNETOM Prismafit 3-Tesla scanner equipped with a 20-channel phased array coil. The children were positioned supine on the scanner bed and presented with food cue stimuli through a mirror attached to the head coil. Blood-oxygen-level-dependent (BOLD) functional scans were obtained using a single-shot gradient echo planar imaging sequence with these parameters: repetition time (TR) of 2000 milliseconds (ms), echo time (TE) of 25 ms, bandwidth of 2520 Hz/pixel, flip angle of 85°, field of view of 220 × 220 mm, matrix size of 64 × 64, and a slice thickness of 4 mm, resulting in 32 slices covering the entire brain. A 3D Magnetization Prepared Rapid Gradient Echo (MPRAGE) sequence was also collected as a structural template for multi-subject registration. Here, parameters were: repetition time of 1950 ms, echo time of 2.26 ms, flip angle of 9°, inversion time of 900 ms, matrix size of 256 × 256 × 224, and a voxel resolution of 1 × 1 × 1 mm^3^. The functional scan lasted for 196 seconds, and the structural scan for 4 minutes and 14 seconds.

### 2.4. Image preprocessing

The whole analysis (preprocessing, first-level, second-level, functional connectivity) of the fMRI data was conducted using Statistical Parametric Mapping (SPM12, http://www.fil.ion.ucl.ac.uk/spm/software/spm12) software, which is implemented in Matlab (MathWorks, Natick, MA, USA). Prior to analysis, the functional images underwent slice time correction and realignment. The anatomical images were then co-registered to the mean functional image and segmented using unified segmentation, which allowed for normalization of the structural image to the standard Montreal Neurological Institute (MNI) space. The resulting transformation matrix was then used to normalize the realigned functional images. Prior studies in children used a standard MNI template for transformation (Bohon, 2017; Boutelle et al., 2015; Luo et al., 2019). Finally, the functional images were smoothed with an 8mm FWHM Gaussian kernel. For noise correction, white matter and cerebrospinal fluid (CSF) signals were extracted from the normalized functional images using the PhysIO toolbox (Kasper et al., 2017).

Participants were excluded from the analyses if they exhibited excessive motion (threshold of 2mm or 2° in any direction) in at least one visit (adapted to (Luo et al., 2019). Only participants with accepted recordings before and after glucose ingestion were included. In addition, a visual inspection was also conducted to identify any potential artefacts. In one case, the exclusion was based on a metallic artefact. In the other case, it was due to the presence of brain lesions. Out of the initial 112 participants recruited, 51 participants were excluded due to excessive movement exceeding 2mm or 2° in any direction in at least one visit or to artefacts during acquisition. Eight additional participants were excluded because they did not finish the oGTT. This resulted in 53 participants who were included for all analyses.

### 2.5. Neural food cue reactivity

For the first-level analyses, brain responses to stimuli were modelled for each participant as blocks convolved with a canonical hemodynamic response function using the general linear model. Two regressors representing high-caloric food images (F) and non-food images (NF) were included, respectively. To account for head motion, the six realignment parameters were included as confounds. The PCA based extracted white matter and CSF signals were also added as confounds. The data were high-pass filtered with a cut-off of 128s and separate contrast images between F versus NF were calculated before and after glucose drink for each individual. In the second-level analysis, the F-NF images obtained from the first-level analysis were used in a full-factorial model (described below, see 2.7).

### 2.6. Functional connectivity. Generalized psychophysiological interaction (gPPI)

To investigate FC during the food cue task, a generalized psychophysiological interaction (gPPI) analysis was employed ((McLaren, Ries, Xu, & Johnson, 2012), https://www.nitrc.org/projects/gppi, version 13.1), which is a well established method for FC analysis in task-based fMRI.

Nine seeds were defined based on significant neural food cue reactivity (F-NF contrast), using a spherical ROI with 6mm (p<0.05, FWE-cluster level; see *Table 2*).

Functional brain connectivity was then calculated for each of these seed regions. The gPPI contrast images of F vs. NF for each seed region (and visit) were included in a second-level full-factorial model (described below, see 2.7).

### 2.7. Second level statistics

In the second-level analysis, the F-NF images (neural food cue reactivity and gPPI) obtained from the first-level analysis were used in a full-factorial model with ISI and sex as between-subject factors, and before vs. after glucose ingestion as a within-subject factor (time-point). The assignment of children with high or low ISI was based on median split (ISI-Matsuda range: 0.94 – 26.57, median split at 8.96). Age and BMIz were included as adjusted covariates. The primary aim of the analysis was to evaluate the main effect of ISI (low vs high) and before and after glucose ingestion (time-point) on neural food-cue reactivity and functional connectivity. In exploratory analyses, interactions between sex and ISI and time-point were evaluated. Statistical significance was determined using a primary threshold of p<0.001 uncorrected and a secondary threshold of p<0.05 family wise error corrected for multiple comparisons at the cluster level (cFWE). We used the SPM Cluster Threshold toolbox (https://github.com/CyclotronResearchCentre/SPM_ClusterSizeThreshold) to calculate the minimum number of voxels which determine a significant cluster. In addition, small volume correction (svc) was performed for insulin-sensitive brain regions-of-interest (ROI’s) including the bilateral hypothalamus, nucleus accumbens, caudate nucleus, putamen, pallidum, hippocampus, amygdala, insula, and PFC (Kullmann et al., 2020). These regions were combined in one single mask. The masks were based on the AAL atlas 3 (https://www.oxcns.org) and wfu_PickAtlas (https://www.nitrc.org/projects/wfu_pickatlas/)

## 3. Results

### 3.1. Participants

The 53 children included 29 girls and 24 boys, with a mean and standard deviation (SD) age of 8.59±0.99 years. The median ISI based Matsuda index was 8.96. *Table 1* presents the comparison of sex, age, adiposity measures, HOMA IR and Matsuda ISI for the low ISI vs. high ISI group. Boys and girls are similarly distributed across the groups. As expected, high ISI is associated with lower adiposity and insulin resistance. *Supplementary table 1* provides additional information on the general characteristics of the participants.

**Table 1.**
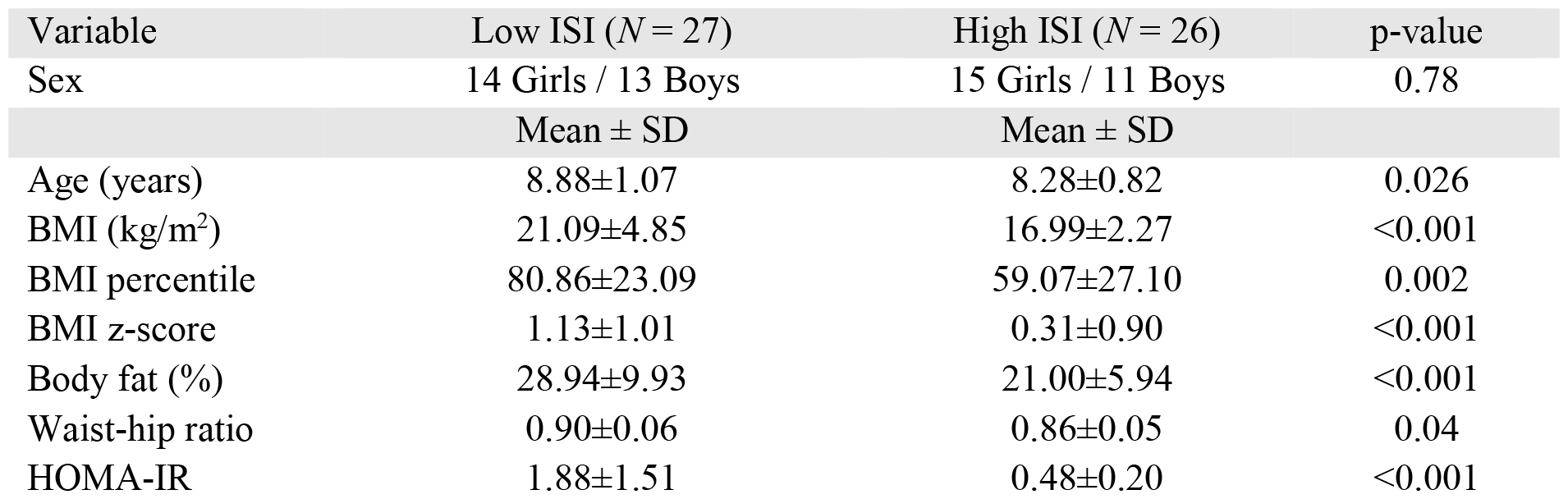
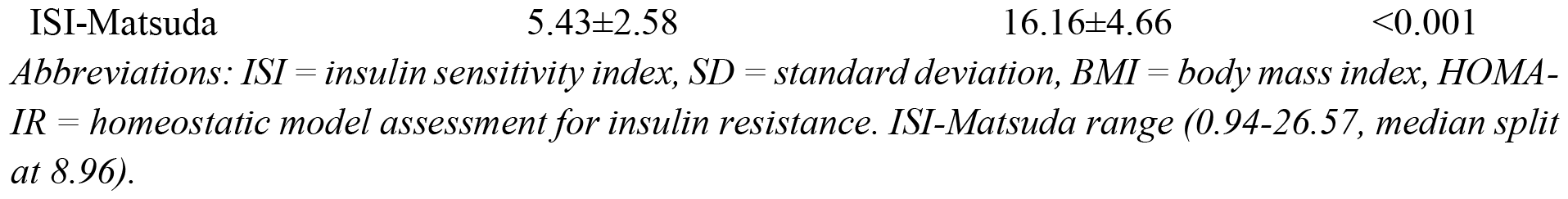
Participants’ characteristics.

**Table 2.** Brain food cue response (Food > non food)

| Brain region | Peak MNI coordinates<br>(X,Y,Z) | Peak t | p(cFWE) | Cluster<br>size |
| --- | --- | --- | --- | --- |
| R posterior Insula | 39, -4, 8 | 7.19 | 0.012 | 116 |
| R anterior Insula | 36, 5, -13 | 5.20 |  |  |
| L posterior Insula | -39, -7, 8 | 7.12 | <0.001 | 237 |
| L anterior Insula | -36, 2, -10 | 5.58 |  |  |
| R Lateral occipital cortex | 45, -64, -4 | 6.21 | 0.018 | 104 |
| L Lateral occipital cortex | -45, -64, 4 | 6.16 | <0.001 | 241 |
| R Postcentral gyrus | 60, -19, 41 | 6.13 | <0.001 | 462 |
| L Postcentral gyrus | -57, -28, 47 | 5.67 | <0.001 | 349 |
| L Amygdala | -15, -4, -22 | 6.07 | 0.031 | 89 |

Table presents areas in which activation was higher following presentation of high-caloric food compared to non-food pictures, before and after glucose ingestion. Labels from the SPM Neuromorphometrics atlas, adapted for children. Abbreviations: L = Left, R = Right, cFWE = family wise error corrected for multiple comparisons at the cluster level.

### 3.2. Neural food cue reactivity

In the whole brain analysis, we observed several regions that were more activated during visual presentation of high-caloric food images compared to non-food images (F-NF contrast, *Figure 1, Table 2*). This corresponds to findings of previous studies investigating neural food cue responsivity (Luo et al., 2019). The coordinates reported in *Table 2* correspond to the centre of the 9 peak coordinates extracted and used as seed regions for the gPPI analysis (6 mm spherical ROIs). No significant main effects of time-point (before and after glucose ingestion), sex, or ISI were found. No significant 2-way interactions were found.

**Figure 1.**
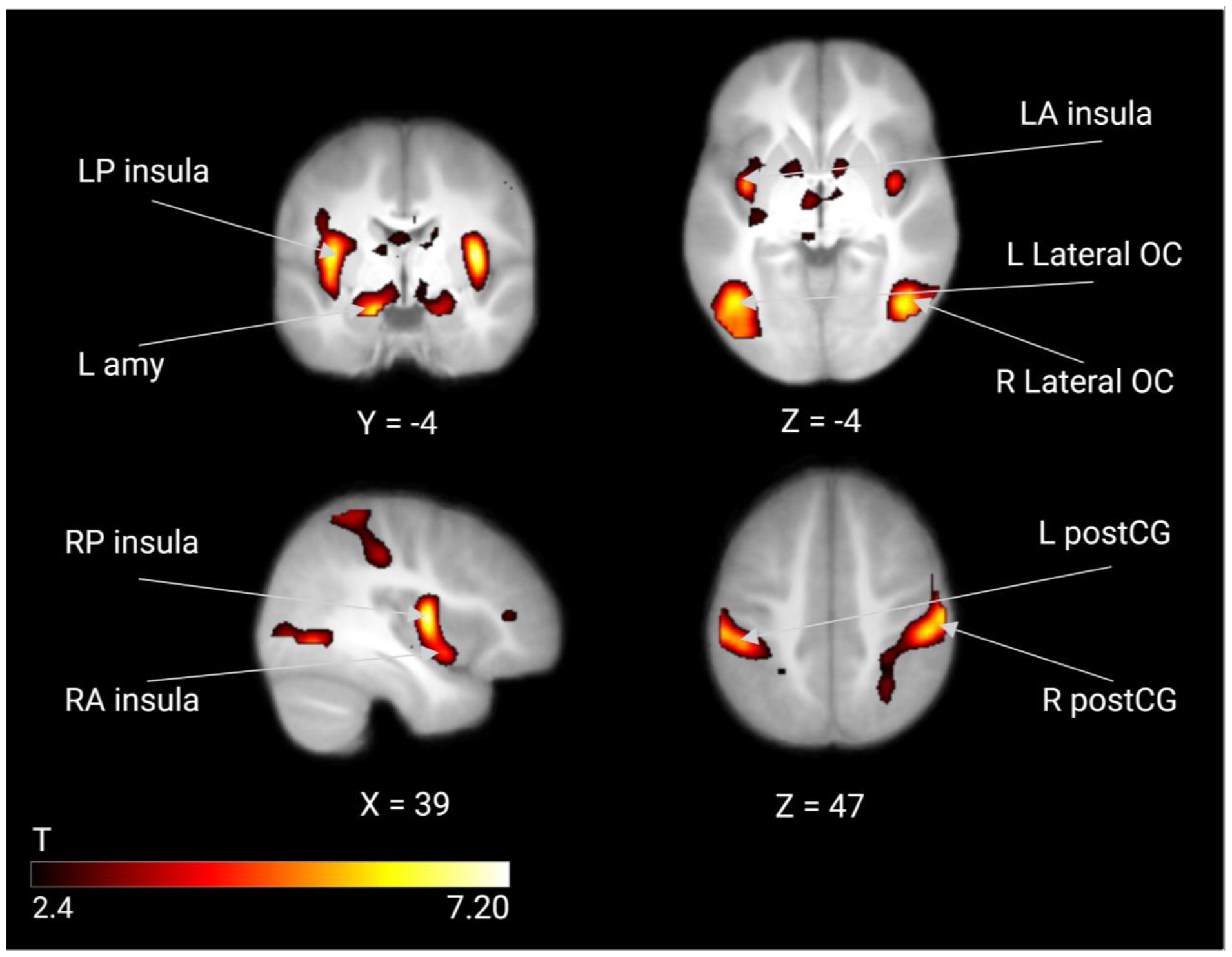
Neural food cue reactivity. Figure shows food cue responsive brain areas based on the F-NF contrast. These regions were used as seeds for gPPI functional connectivity analysis. Color map corresponds to T values (P < 0.001 uncorrected for display) overlaid on the average normalized T1 weighted image of the children. R = right; L = left; P = posterior; A = anterior; amy = amygdala; postCG = postcentral gyrus; OC, occipital cortex.

### 3.3. Task-based functional connectivity analysis

We investigated functional connectivity in response to food cues before and after glucose ingestion (time-point) in children with high and low peripheral insulin sensitivity. There was a significant main effect based on ISI group such that FC was higher between the left anterior insula and the right nucleus caudate in the low ISI compared to the high ISI group (*Figure 2, Table 3*). In addition, we found lower FC between the left posterior insula and the right middle temporal gyrus (MTG) in the low ISI compared to the high ISI group (*Figure 3, Table 3*).

**Table 3.**
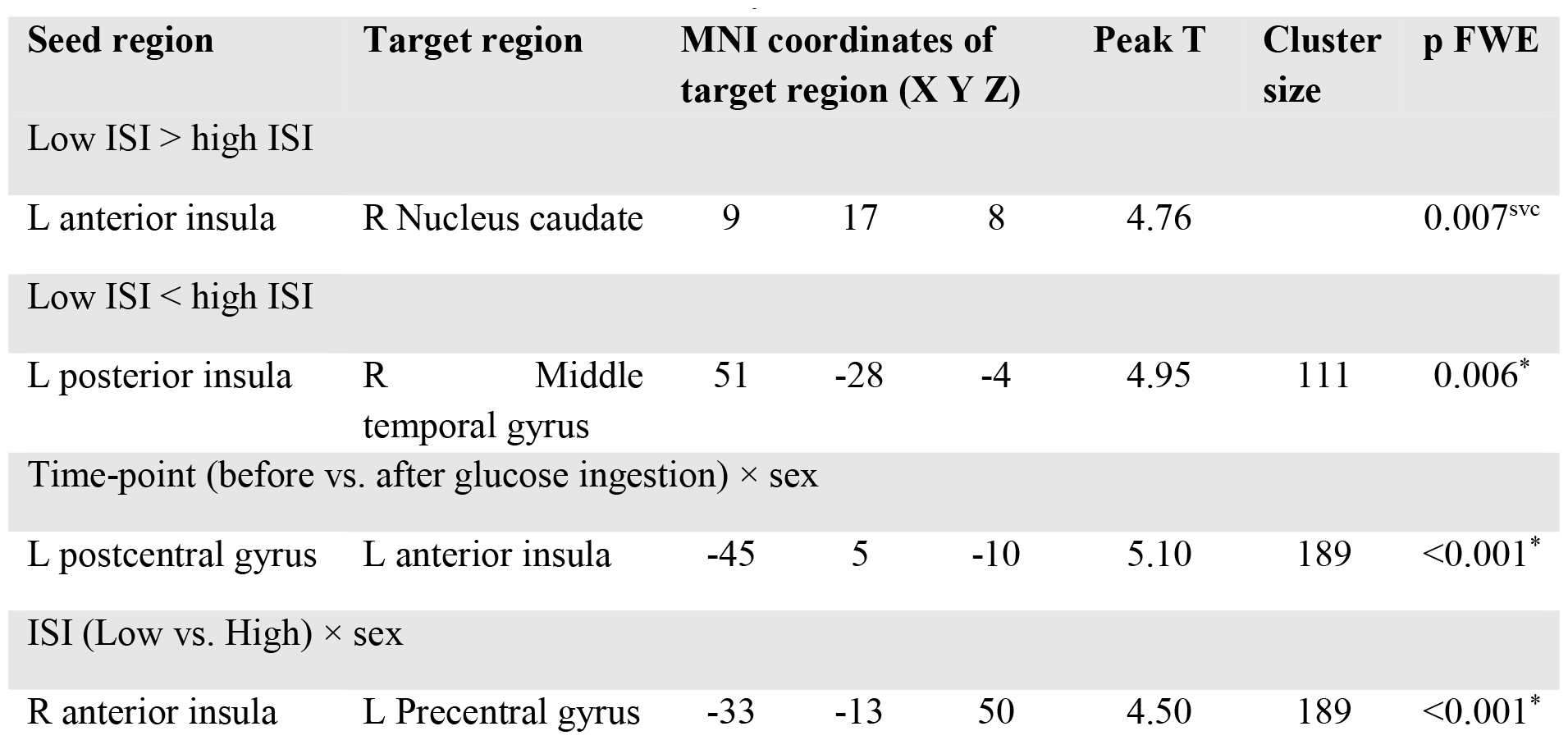
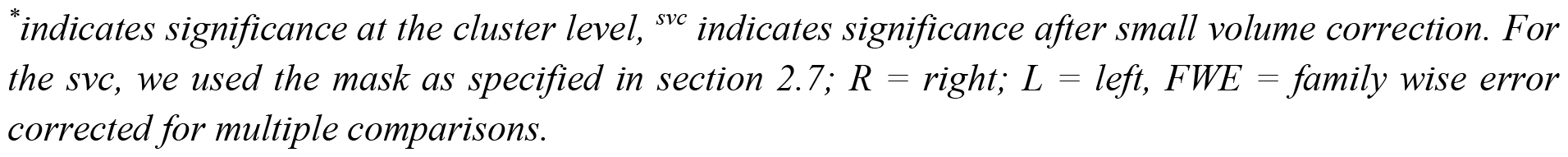
Food-cue induced functional connectivity (food vs. non-food)

| Seed region | Target region | MNI coordinates of target region (X Y Z) |  |  | Peak T | Cluster size | p FWE |
| --- | --- | --- | --- | --- | --- | --- | --- |
| Low ISI > high ISI |  |  |  |  |  |  |  |
| L anterior insula | R Nucleus caudate | 9 | 17 | 8 | 4.76 |  | 0.007<sup>svc</sup> |
| Low ISI < high ISI |  |  |  |  |  |  |  |
| L posterior insula | R Middle temporal gyrus | 51 | -28 | -4 | 4.95 | 111 | 0.006<sup>*</sup> |
| Time-point (before vs. after glucose ingestion) × sex |  |  |  |  |  |  |  |
| L postcentral gyrus | L anterior insula | -45 | 5 | -10 | 5.10 | 189 | <0.001<sup>*</sup> |
| ISI (Low vs. High) × sex |  |  |  |  |  |  |  |
| R anterior insula | L Precentral gyrus | -33 | -13 | 50 | 4.50 | 189 | <0.001<sup>*</sup> |

\* indicates significance at the cluster level, <sup>svc</sup> indicates significance after small volume correction. For the svc, we used the mask as specified in section 2.7; R = right; L = left, FWE = family wise error corrected for multiple comparisons.

**Figure 2.**
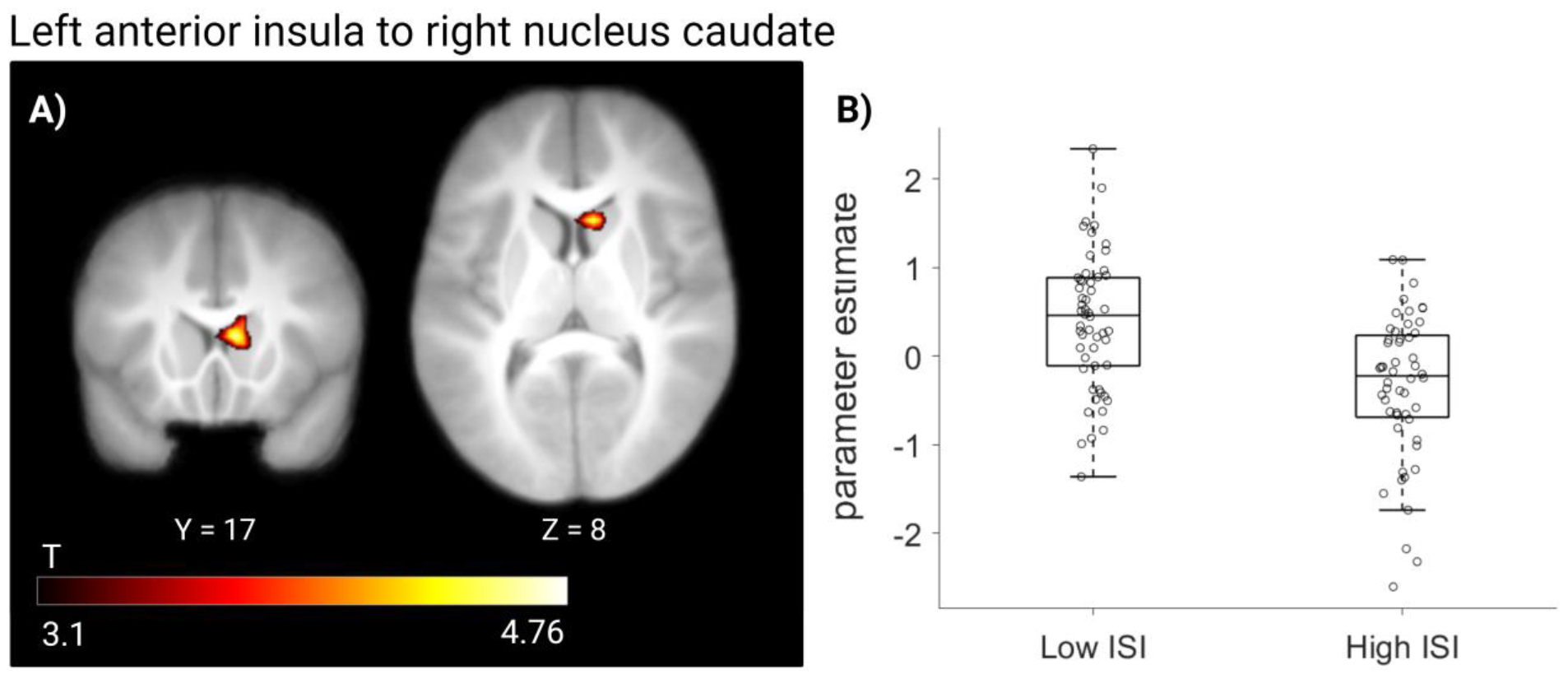
Higher food-cue induced functional connectivity in low ISI compared to high ISI children. A) Shown is the cluster of the right nucleus caudate, revealing higher FC with the anterior insula in children with lower peripheral insulin sensitivity. Color map corresponds to T values (P < 0.001 uncorrected for display) overlaid on the average normalized T1 weighted image of the children. B) Box plots show the left anterior insula FC to the right nucleus caudate in children with lower and higher peripheral insulin sensitivity (low ISI and high ISI, respectively). Whiskers indicate 1.5 interquartile range. ISI = Insulin Sensitivity Index

**Figure 3.**
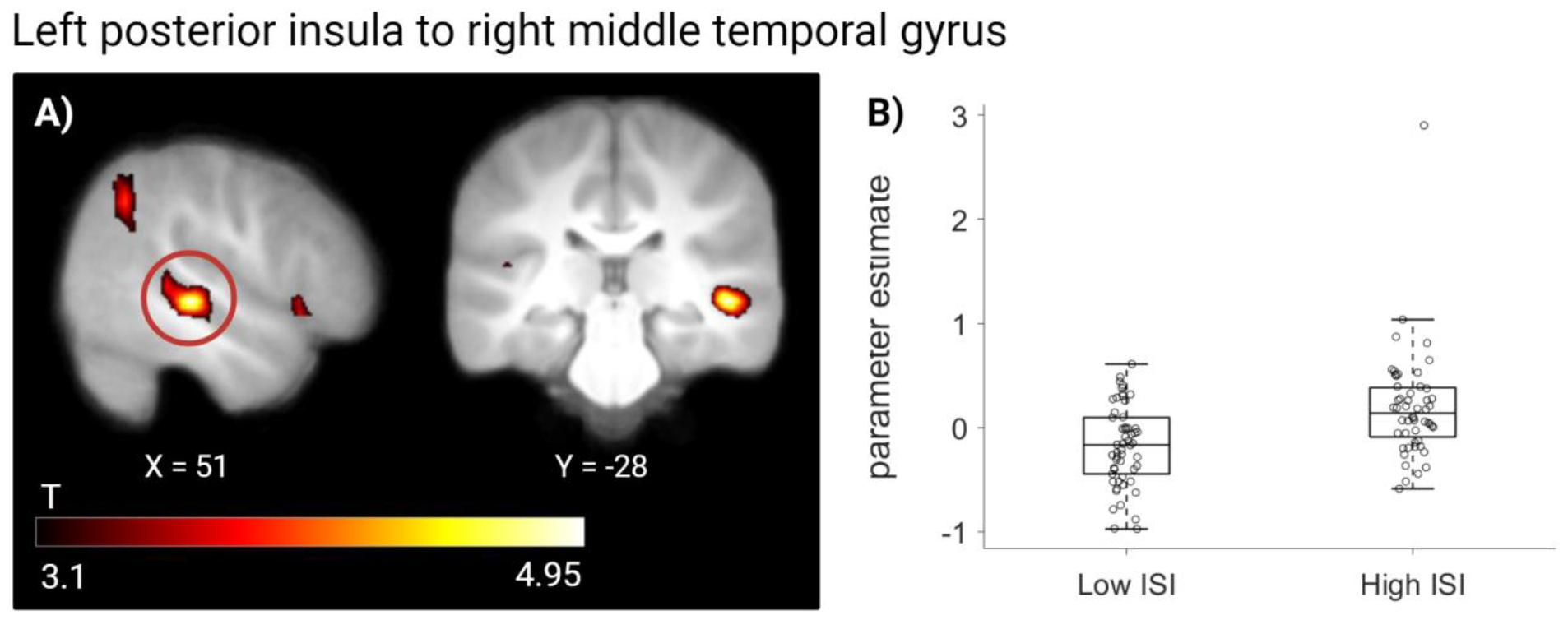
Lower food-cue induced functional connectivity in low ISI compared to high ISI children. A) Shown is the cluster in the middle temporal gyrus, revealing lower FC with the left posterior insula in children with lower peripheral insulin sensitivity. Colour maps on the left correspond to T values (P < 0.001 uncorrected for display) overlaid on the average normalized T1 weighted image of the children. B) Box plots show the left posterior insula FC to the right middle temporal gyrus in children with lower and higher peripheral insulin sensitivity (low ISI and high ISI, respectively). Whiskers indicate 1.5 interquartile range. ISI = Insulin Sensitivity Index

We found an interaction between time-point × sex in FC between the left postcentral gyrus and the left anterior insula (*Figure 4, Table 3*). FC in boys increased from before to after glucose ingestion, while it decreased in girls (Post hoc tests: Boys before < Boys after: t(46) = −2.60, p ≤ 0.01; Girls before > Girls after: t(56) = 4.05, p ≤ 0.001; Boys before < Girls before: t(51) = −2.70, p ≤ 0.01; Boys after > Girls after: t(51) = 3.61, p ≤ 0.001). We found an interaction between ISI × sex in FC between the right anterior insula and the left precentral gyrus (*Figure 5, Table 3*). Boys with low ISI and girls with high ISI showed the strongest FC (Post hoc test: Boys low ISI > Boys high ISI: t(46) = 4.28, p ≤ 0.0001; Girls low ISI < Girls high ISI: t(56) = −3.61, p ≤ 0.001; Boys low ISI > Girls low ISI: t(52) = 3.70, p ≤ 0.001; Boys high ISI < Girls high ISI: t(50) = −4.34, p ≤ 0.0001). No main effects of sex or of time-point were found. No other interactions were detected.

**Figure 4.**
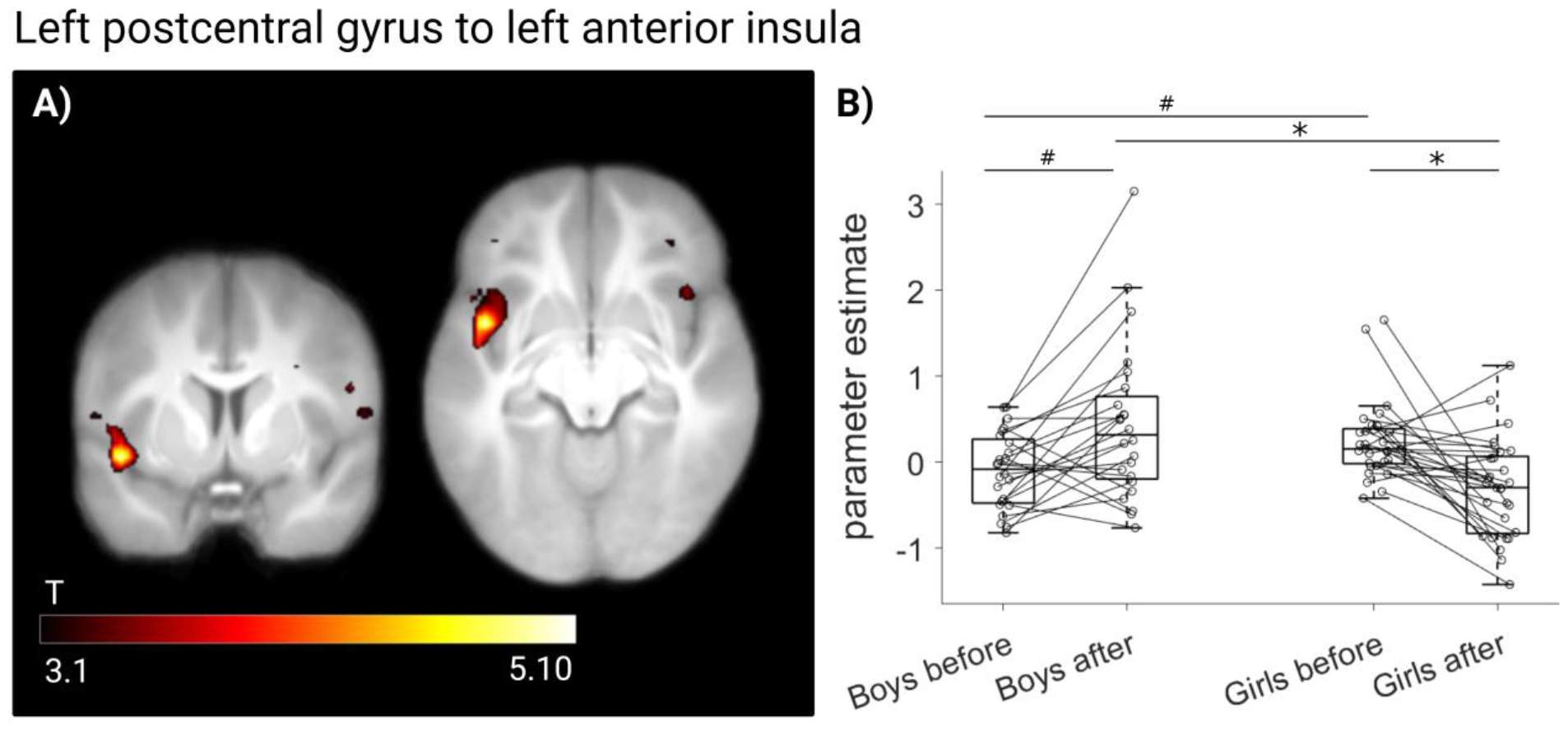
Interaction between time-point (before and after glucose ingestion) and sex on functional connectivity in children. A) Shown are the clusters of the left postcentral gyrus network, revealing an interaction between sex and time-point (before and after glucose ingestion). Colour maps on the left correspond to T values (P < 0.001 uncorrected for display) overlaid on the average normalized T1 weighted image of the children. B) Box plots showing the left postcentral gyrus FC to the left anterior insula in boys and girls before and after glucose ingestion. Whiskers indicate 1.5 interquartile range. ^*^ indicates p≤0.001, ^#^ p≤0.01.

**Figure 5.**
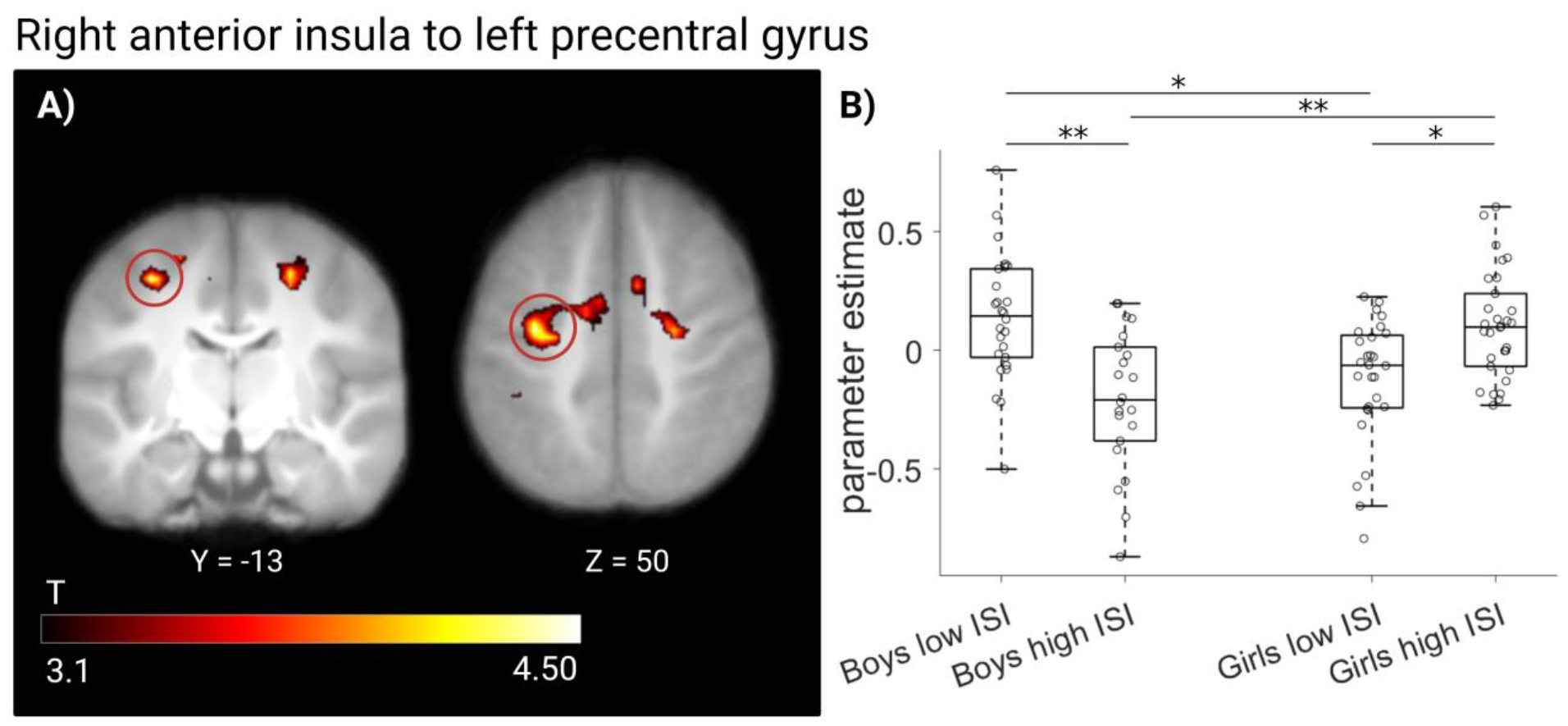
Interaction between peripheral insulin sensitivity and sex on functional connectivity in children. A) Shown are the clusters of the right anterior insula network, revealing an interaction between sex and peripheral insulin sensitivity. Colour maps on the left correspond to T values (P < 0.001 uncorrected for display) overlaid on the average normalized T1 weighted image of the children. B) Box plots showing the right anterior insula FC to the left precentral gyrus in boys and girls with lower and higher peripheral insulin sensitivity (low ISI and high ISI, respectively). Whiskers indicate 1.5 interquartile range. ^*^ indicates p≤0.001, ^**^ p≤0.0001. ISI = Insulin Sensitivity Index

## 4. Discussion

The aim of this study was to examine the relationship between peripheral insulin sensitivity and neural food-cue induced activity and functional connectivity (FC) both before and after glucose ingestion in children aged between 7 and 11 years from the BrainChild Cohort (Alves et al., 2020; Luo et al., 2019; Luo et al., 2021; Lynch et al., 2021). Neural food cue reactivity reported in the current study corresponds to previous works in the BrainChild study (Luo et al., 2019), with higher activity to food compared to non-food in the bilateral posterior and anterior insula, lateral occipital cortex, postcentral gyrus, and left amygdala to food compared to non-food visual cues. We found no significant effects of peripheral insulin sensitivity on whole-brain food cue reactivity. However, we observed different insular FC patterns in response to high caloric food-cues in children with higher compared to lower insulin sensitivity in the fasted state and after glucose ingestion. Children with lower insulin sensitivity had higher insular connectivity towards food-reward regions, and lower connectivity towards cognitive control regions. In an explorative analysis, we found sex differences in food-cue functional connectivity. Our results suggest that children with lower peripheral insulin sensitivity, independent of body mass index, have alterations in FC between food-cue responsive brain regions regardless of prandial state.

### 4.1. Insular functional connectivity during food cues is associated with ISI

While we found differences in FC during food cue task with respect to peripheral insulin sensitivity, we did not observe differences in whole brain reactivity to food images. This corresponds to a recent meta-analysis showing that there is little evidence for obesity related differences on whole brain food cue reactivity in children and adults suggesting that there are other mediating factors that may not have been considered thus far (Morys, García-García, & Dagher, 2020). Our results show, however, distinct patterns of insular FC during the viewing of food-cues in children based on their peripheral insulin sensitivity. Generally, reduced insulin responsiveness in the insula cortex has been observed in adults with peripheral insulin resistance and poor cognitive control (Wagner et al., 2022; Wagner et al., 2023). In our work, we found differences in connectivity independent of prandial state in two regions of the insula, an anterior and a posterior region that are known to underlie brain responses to taste and visual presentation of food (Avery et al., 2020; Avery, Liu, Ingeholm, Gotts, & Martin, 2021). However, these two insular regions also present functional differences. While the dorsal, posterior region is associated with cognitive and attentional processing, the anterior ventral insula is primarily involved in social and emotional functions. Connections of the two insular regions are also different. Specifically, the posterior section is mainly connected with frontoparietal regions, while the anterior ventral region is connected with emotional regions in the limbic system. For more details see (Avery et al., 2021; Kurth, Zilles, Fox, Laird, & Eickhoff, 2010). In agreement with our hypothesis, we observed higher FC between areas responsive to high-caloric food images in children with low ISI vs. high ISI, with higher FC between the left anterior insula and the right caudate (*Figure 2*). The insula and caudate play a crucial role in associating food cues with rewards, in wanting and craving, and attributing subjective value to palatable food (Kahathuduwa, Boyd, Davis, O’Boyle, & Binks, 2016). Hyper- or hypoactivations in neural activity along these areas is associated with alterations in reward processing, inhibitory control, and body weight regulation (Li et al., 2023a) and may lead to overeating due to increased sensitivity to food cues (Meng, Huang, Ao, Wang, & Gao, 2020; Rothemund et al., 2007). Prior studies have shown that adults with obesity (Stoeckel et al., 2009; Wijngaarden et al., 2015) and lean adolescents at high risk of developing obesity (Sadler et al., 2023) have higher FC between reward-related areas, which is thought to contribute to the elevated motivational value of food in this population. Specifically, heightened connectivity between the anterior insula and caudate was previously interpreted as one factor that may lead to overeating by impairing self-awareness, increasing arousal in response to food cues, and reducing responsiveness to a post-prandial state (Donofry et al., 2020b; Geha, Cecchi, Todd Constable, Abdallah, & Small, 2017; Nummenmaa et al., 2012). In light of these previous reports, the heightened connectivity between anterior insula and caudate in children with lower peripheral insulin sensitivity might be an early marker of insulin resistance and risk for the development of obesity in youth.

In addition, we found that children with low insulin sensitivity have lower FC between the left posterior insula and the right middle temporal gyrus (MTG, *Figure 3*). The MTG is involved in emotional memory and processing of food odors (Dolcos, LaBar, & Cabeza, 2005; Han, Roitzsch, Horstmann, Pössel, & Hummel, 2021; Kohn et al., 2014). Adults with obesity exhibit reduced intrinisic activity in the MTG following intermittent energy restriction (Li et al., 2023b). The authors attributed this reduction to a potential decrease in cognitive functions and neural processing of sensory information, which could in turn impact eating behavior. The role of MTG in obesity is found also in earlier life stages. For example, a lower FC in the MTG was found in adolescents with overweight and obesity (Moreno-Lopez, Contreras-Rodriguez, Soriano-Mas, Stamatakis, & Verdejo-Garcia, 2016). Furthermore, in children aged 7 to 9 with obesity, there was a lower brain response in the MTG when exposed to high-energy food pictures (Masterson et al., 2019). These results are consistent with the idea that metabolic conditions like overweight, obesity, and insulin resistance may have an effect not only on the brain networks responsible for metabolic control and reward, but also on more complex cognitive functions (Moreno-Lopez et al., 2016; Verdejo-García et al., 2010).

### 4.2. Sex differences on food cue induced FC

No overall sex differences were found on food-cue reactivity and FC. However, sex differences in FC were found depending on prandial state and on peripheral insulin sensitivity. Specifically, FC between the left postcentral gyrus and the left anterior insula increased after glucose ingestion in boys, and decreased in girls (*Figure 4*). In addition, when comparing children with different ISI levels, we observed that boys with lower ISI had higher FC, whereas this trend was reversed in girls (*Figure 5*). Generally, sex effects on insula and sensorimotor regions functional connectivity have been reported before in the context of obesity and metabolic research. However, the outcomes of these studies do not yet show a clear pattern. For example, there is some indications that men (vs. women) with obesity had higher FC in response to food cues with the amygdala in supplementary and primary motor areas (Atalayer et al., 2014; Kilpatrick, An, Pawar, Sood, & Gupta, 2023). In addition, men showed a more prominent decrease in postprandial insular resting-state connectivity with sensorimotor and prefrontal cortex (Kilpatrick et al., 2020). Based on these results, the authors have suggested an increased vulnerability of males to obesity-related alterations in the precentral gyrus and occipital cortex (Gupta et al., 2017; Kilpatrick et al., 2023). In general, however, alterations in somatosensory regions in children with overweight and obesity might indicate an expanded somatosensory and motor cortical representation of the body as a function of body mass (Pujol et al., 2021). In their work, the authors found that higher body mass was associated with higher integration of the sensorimotor cortex to superior parietal regions that underlie body awareness. Possible sex differences might also depend on the different subcutaneous and visceral fat distribution between males and females which are ultimately related to insulin resistance (Machann et al., 2005).

## 5. Limitations

In the current analysis we recognize some limitations. The cross-sectional nature of the analysis precludes inference about the directionality between children’s peripheral insulin sensitivity and the differences in brain functional connectivity. In addition, the ∼15 min interval between the two food cues tasks (before and after glucose ingestion) may not capture neural food cue processing when circulating insulin levels are at their peak. Finally, in our analysis we focused on task-related FC during a food cue task. Analysis of resting-state connectivity is also useful to study brain network changes at a young age (Brooks et al., 2023).

## 6. Conclusion

The study investigated the relationship between peripheral insulin sensitivity and both, neural reactivity and whole brain FC in children during the processing of high-caloric food cues. Based on our findings, we found no relation between peripheral insulin sensitivity and whole-brain food reactivity. However, there was an association between low peripheral insulin sensitivity and insular functional connectivity between food processing, reward, and cognitive regions in the brain. In addition, sex-specific patterns of functional connectivity were depending on both peripheral insulin sensitivity and prandial state (fasted vs. after glucose ingestion). Overall, the results support the idea that insulin resistance, independent of adiposity, can alter communication along brain regions related to food processing, reward, regulation of food intake, and emotions. These findings are consistent with previous research on obesity and suggest that similar pathways may be involved in the development of obesity and insulin resistance. These findings align with the notion that metabolic conditions such as overweight, obesity, and insulin resistance impact the brain networks responsible for metabolic regulation and reward and influence higher-order cognitive processes.

## 7. Acknowledgements

This work was supported by an American Diabetes Association Pathway Accelerator Award (#1-14-ACE-36) (PI: K.A.P) and the National Institute of Diabetes and Digestive and Kidney Diseases (NIDDK), National Institutes of Health (NIH) R03DK103083 (PI: K.A.P), R01DK116858 (PIs: K.A.P, A.H.X). A Research Electronic Data Capture, REDCap, database was used for this study, which is supported by the Southern California Clinical and Translational Science Institute (SC CTSI) through NIH UL1TR001855. P.M.T. is funded in part by NIH grant U54 EB020403. The work of L.S, H.P. and S.K. was supported by a grant (01GI0925) from the Federal Ministry of Education and Research (BMBF) to the German Center for Diabetes Research (DZD e.V.). We also thank the International Max Planck Research School for the Mechanisms of Mental Function and Dysfunction (IMPRS-MMFD) and the Add-on Fellowship of the Joachim Herz Foundation for the support to L.S.

## 8. Author contributions

L.S conceptualized and conducted the analysis, drafted and revised the manuscript, and prepared the figures. R.V. supported the analysis and supervised the work. S.Z. supported the analysis and discussed the results. A.H.X., S.L. and K.A.P. conceptualized the original study, performed the recordings, have full access to all data in the study and take responsibility for the integrity of the data. B.A. performed the recordings. A.L.B., H.P., S.K. supervised the work. All authors discussed the results and implications, reviewed and edited the manuscript and approved its final version.

