## Supplementary table 1 for "Influence of insulin sensitivity on food cue evoked functional brain connectivity in children"

*Supplementary table 1.* General participants characteristics

| Variable | Mean ± SD | Range |
|---|---|---|
| Age (years) | 8.56 ± 0.95 | 7.33 – 11.23 |
| BMI (kg/m$^2$) | 18.79 ± 4.25 | 12.11 ± 34-01 |
| BMI percentile | 67.85 ± 29.07 | 0.06 ± 99.58 |
| BMI z-score | 0.77 ± 1.05 | -1.64 ± 2.64 |
| Body fat (%) | 25.05 ± 9.07 | 12.7 ± 57.10 |
| Waist – hip ratio | 0.88 ± 0.05 | 0.79 ± 1.04 |
| Insulin resistance (Homa-IR) | 1.19 ± 1.28 | 0.23 ± 7.58 |
| Insulin sensitivity (ISI-Matsuda) | 10.69 ± 6.56 | 0.94 ± 26.57 |
| Sex | 29 females 24 males | |
| Pubertal Tanner | Tanner1=48; Tanner2=3; Tanner3=2 | |
